# Increasing computational protein design literacy through cohort-based learning for undergraduate students

**DOI:** 10.1101/2022.05.24.493341

**Authors:** Erin C. Yang, Robby Divine, Christine S. Kang, Sidney Chan, Elijah Arenas, Zoe Subol, Peter Tinker, Hayden Manninen, Alicia Feichtenbiner, Talal Mustafa, Julia Hallowell, Isiac Orr, Hugh Haddox, Brian Koepnick, Jacob O’Connor, Ian C. Haydon, Karla-Luise Herpoldt, Kandise Van Wormer, Celine Abell, David Baker, Alena Khmelinskaia, Neil P. King

**Author notes:** these authors contributed equally (Erin C. Yang, Robby Divine). Correspondence should be addressed to A.K. and N.P.K.

## Abstract

Undergraduate research experiences can improve student success in graduate education and STEM careers. During the COVID-19 pandemic, undergraduate researchers at our institution and many others lost their work-study research positions due to interruption of in-person research activities. This imposed a financial burden on the students and eliminated an important learning opportunity. To address these challenges, we created a paid, fully-remote, cohort-based research curriculum in computational protein design. Our curriculum used existing protein design methods as a platform to first educate and train undergraduate students and then to test research hypotheses. In the first phase, students learned computational methods to assess the stability of designed protein assemblies. In the second phase, students used a larger dataset to identify factors that could improve the accuracy of current protein design algorithms. This cohort-based program created valuable new research opportunities for undergraduates at our institute and enhanced the undergraduates’ feeling of connection with the lab. Students learned transferable and useful skills such as literature review, programming basics, data analysis, hypothesis testing, and scientific communication. Our program provides a model of structured computational research training opportunities for undergraduate researchers in any field for organizations looking to expand educational access.

**Graphical Abstract:** 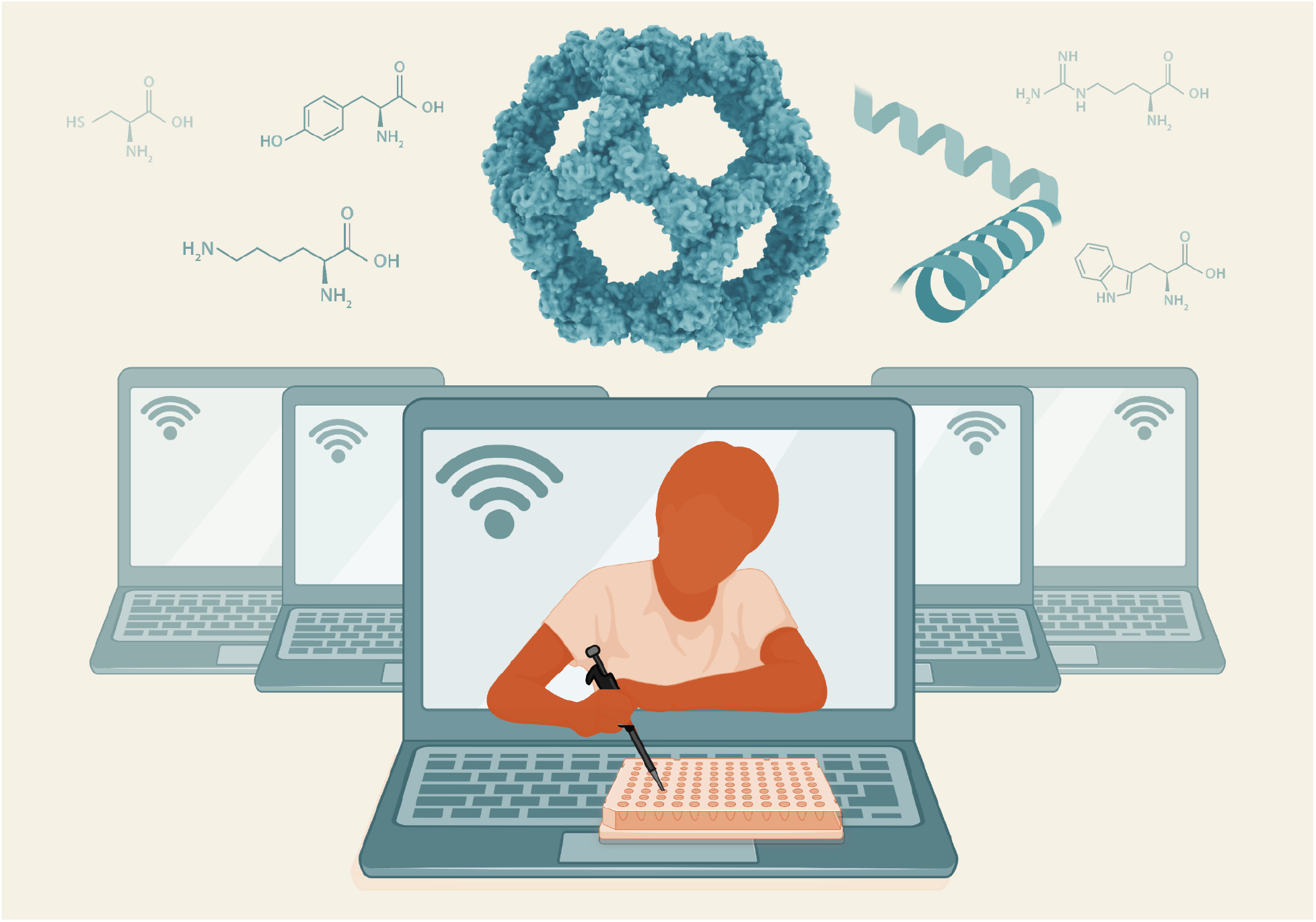

## Introduction

Participation in hands-on research fuels students’ interest in pursuing STEM graduate education and careers^1–3^ and also heightens graduate education performance in relevant research skills.^4^ Several studies have correlated research experiences with increased retention,^5^ improved academic performance,^6^ and persistence in STEM courses.^7^ Through undergraduate research experiences, students are exposed to laboratory techniques, analytical thinking, autonomous problem solving, and collaboration. Students’ motivation to learn is also enhanced by research experiences.^8–10^ Exposure to non-classroom science, guidance from mentors in the lab environment, and peer relationships during their research experience are particularly valuable for students learning about STEM careers at minority-serving institutions.^11^ However, all of these benefits are limited by mentor and resource availability.

Prior to the COVID-19 pandemic, undergraduate research at the Institute for Protein Design at the University of Washington was based on an apprenticeship model.^1,10,12–14^ Undergraduates spent two quarters learning about the organization of the Institute and providing general lab support to researchers. After these initial two quarters, undergraduates were granted the opportunity to conduct an independent research project under the mentorship of a graduate student or postdoctoral researcher at the Institute. Projects ranged from solely computational to solely experimental, depending on the project scope decided between the mentor and trainee. Each project was tailored to the undergraduate, who was encouraged to present their work at a university-wide Undergraduate Research Symposium held at the end of each academic year.^15^ While the apprenticeship model is a well-established approach, it is limited by mentor availability, research ideas with an appropriate scope for an undergraduate research project, and the mentee’s graduation timeline. The apprenticeship model is particularly difficult at institutions where research is not central to the institutional mission, especially for students who do not actively seek out opportunities.^16^ Alternatives to the apprenticeship model that still enable student creativity and independence can help provide diverse, high-quality research opportunities to a wider variety of students.

Due to the COVID-19 pandemic, many undergraduates, including those at the University of Washington, spent nearly a full year learning remotely, and undergraduate researchers lost their on-campus work-study research positions.^17,18^ We set out to re-employ the work-study undergraduate researchers at our Institute by offering a virtual research opportunity. However, we needed a different approach than our previous apprenticeship model, as the frequent 1:1 contact central to that model was impractical to implement remotely and too few mentors were free to individually work with each undergraduate at our Institute. Previous cohort-based research courses in computational protein design, in which multiple students are taught by a smaller number of mentors, enabled them to accommodate greater numbers of students and have shown significant success in increasing students’ enthusiasm for biochemistry and computational biology.^19–21^ We adopted this cohort-based model and curated a completely remote computational curriculum that allowed students to take ownership of their portion of a team research project while learning from each other. We called this program *<u>J</u>obs for <u>U</u>ndergraduates in <u>P</u>andemIc <u>T</u>imes <u>E</u>mergency <u>R</u>esponse* (JUPITER). Through this pilot program, we provided tools for undergraduates to understand basic protein structure concepts, coding conventions, and data analysis, while developing transferable skills such as critical thinking, troubleshooting, collaboration, communication, and motivation. The JUPITER program was designed in two phases: in the first phase, students were taught the concepts behind the protein design process; in the second phase, students reinforced their learning by developing and implementing these concepts. We believe that they will use both the technical and transferable skills they learned from this course in their professional careers.

## Program Overview

Nine undergraduate students participated in JUPITER. These students were majoring in STEM fields at the University of Washington, on track to graduate between the years of 2020 and 2022, and had previously held jobs at the Institute for Protein Design which they lost due to the COVID-19 pandemic and associated University policies. In our cohort-based model (**Figure 1A**), the students learned from each other in a collaborative environment while maintaining parallel research directions.

**Figure 1.**
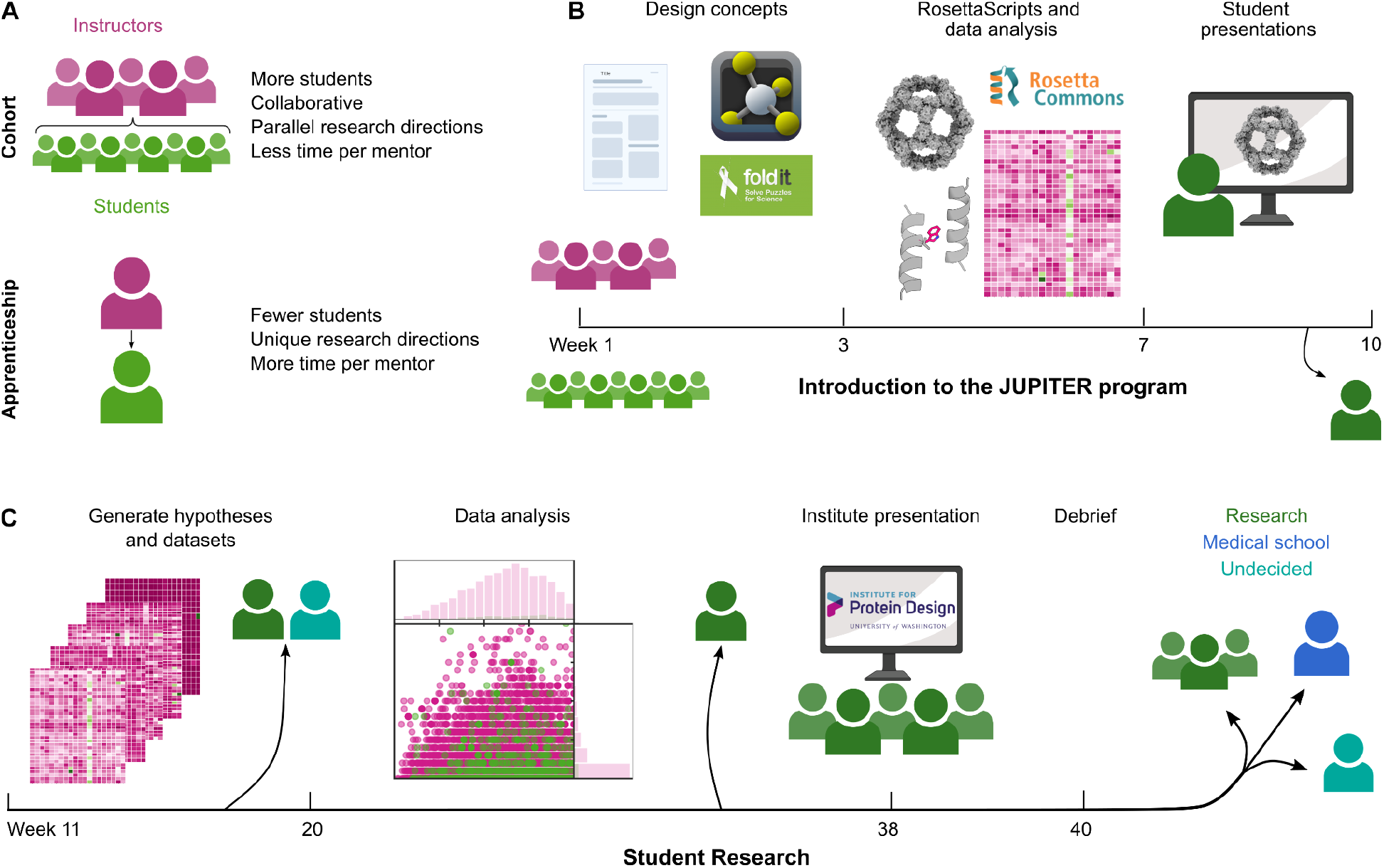
Overview of the JUPITER model and program. **A,** The cohort-based model allows mentors to instruct more students. This model encourages more collaboration between students, compared to a one-on-one apprenticeship model. **B,** In the first 10 weeks of JUPITER, students first learned design concepts and computational approaches. These approaches were then applied to a case study that each student presented to the rest of the cohort. **C,** Over the next several months, students applied their knowledge to generate data on a large design set, proposed their own hypotheses and data processing pipelines required to approach them, and presented their results in an Institute-wide seminar.

We designed the JUPITER curriculum to address an ongoing scientific challenge: improving the success rate of current computational methods for the *de novo* design of self-assembling protein complexes using the Rosetta software suite.^22–34^ These methods focus on docking oligomeric protein building blocks into target symmetric architectures (e.g., icosahedral point group symmetry), followed by protein-protein interface design to generate low-energy interfaces between the building blocks that drive assembly specifically to the target structure. While such assemblies are well-suited for molecular encapsulation or display applications such as cellular delivery and vaccine development,^34–39^ current computational models do not allow us to confidently predict the success or stability of design models. Indeed, many designs aggregate or fail to assemble, due to misfolding or suboptimal assembly conditions.^33,40^ Protein designers often test tens to hundreds of designs to find a handful of nanoparticles that assemble to the desired structure.^34,41,42^

To better understand the pitfalls of current nanoparticle design methods, we aimed to computationally distinguish between successful and unsuccessful protein nanoparticle designs. While several published efforts to stabilize monomeric proteins demonstrated high success rates based on approaches using phylogenetic analysis, structure-based rational design, or sequence-based design,^43–49^ none of these approaches addressed the stability of interfaces between multiple interacting proteins and its effect on design success.^50^ We thus sought to test a recently developed computational pipeline—<u>p</u>rotein <u>S</u>train <u>U</u>nsatisfactoriness and <u>F</u>rustration find<u>ER</u> (pSUFER)—for local sequence optimality evaluation on previously designed protein assemblies of known assembly phenotype.^49,51,52^ pSUFER is a Rosetta software suite-based protocol that estimates the effect of point mutations at a particular position in a protein. We tested the pSUFER protocol on a set of 809 designed protein nanoparticle models in order to: (1) determine the effect of point mutations at the previously designed protein-protein interfaces on free energy; and (2) to identify key residue positions predicted to weaken/strengthen the protein-protein interfaces.

Our teaching and research program had two phases. In the first phase, the students were taught how to read and understand a scientific paper and received an introduction to protein structure and design terminology (week 1);^29–32,53^ were trained to use protein visualization and design software such as PyMOL and FoldIt (weeks 2-3);^54,55^ and practiced computational research approaches including accessing remote computational resources, following Rosetta XML basics, running the pSUFER protocol on a single individually assigned protein nanoparticle,^23,30–32^ and conducting basic data analysis (weeks 4-7). This phase ended with individual presentations to the other program participants (weeks 7-10, **Figure 1B**). The undergraduates met with three regular advisors and occasional guest lecturers via Zoom for a one hour-long lecture and additional optional office hours to troubleshoot homework assignments. Assignments were designed to help the undergraduates learn how to evaluate nanoparticle interfaces over the course of the quarter. Following these initial 10 weeks, one student secured a full-time research position in another computational biology group at the University of Washington, citing the JUPITER program as a key driver of their motivation to pursue research.

In the second phase, we embarked on a longer-term research project with our undergraduate cohort, training them in critical thinking. Students worked together in small teams to generate hypotheses on what influenced predicted assembly phenotype, and in the process generated their own datasets, created data analysis pipelines, and presented their work to an audience of over 90 members of the Institute for Protein Design (**Figure 1C**). A major advantage of the JUPITER phase 2 program was that students could probe different questions regarding the mutational effects of nanoparticle interfaces using a single dataset they collectively generated. The undergraduates used Jupyter Notebooks to write Python scripts for data analyses and visualize their results.^56^ Teams met weekly with two mentors to troubleshoot and uncover alternative analysis methods and global trends across the library of nanoparticles. The students’ final Institute-wide presentations clearly demonstrated their greater understanding of protein structure, Rosetta scripting, Python and Bash coding, and broad scientific thinking. Among the 9 undergraduates that had started in our pilot program, 6 secured full-time research/higher education positions in both computational and wet lab projects, 1 applied to medical school, and 2 decided to pursue alternative opportunities in STEM.

## Phase 1 - Introduction to the pSUFER protocol

After becoming familiar with protein design in general and nanoparticle design concepts specifically, each student applied the pSUFER protocol to one assigned nanoparticle. For this, students used PyMOL to extract the asymmetric unit of their nanoparticle and performed design model minimization with the Rosetta all-atom energy function (**Figure 2A**). Students then applied the FilterScan Mover^57^, which scans all possible mutations at a user-defined residue position and computes the difference in a user-defined metric. More specifically, the students computed the difference in Gibbs free energy (dG) between the wild-type amino acid and every possible point mutant (change in free energy, or ddG) at every residue position they identified as belonging to their nanoparticle interface (**Figure 2B**).^53,55,57^ A decrease in free energy was used to define mutations that were more favorable than wild-type (ddG < 0 Rosetta Energy Units, REU), and an increase in free energy was used to define mutations that were less favorable than wild-type (ddG > 0 REU) (**Figure 2C**). By graphing the calculated ddG for each mutation at each position in a barplot, the students comprehensively identified amino acid identities that were predicted to stabilize or destabilize the protein-protein interface (**Figure 2D**). Residue positions with more than two favorable mutations were deemed “frustrated” (**Figure 2E**), as these positions were more mutable than positions with fewer favorable mutations and thus not yet fully optimized. By contrast, residue positions with up to two favorable mutations were deemed “sensitive” positions. Global analysis of the effects of interface mutations through ddG heatmaps allowed the students to identify specific amino acid identities that were unfavorable at many positions or single positions with many unfavorable mutations (**Figure 2F**). Through phase one, students were taught literature review, programming basics, computational protein analysis, and technical communication, using a model that could easily be extended to teach similar skills in other research contexts. All teaching materials from phase 1 and example scripts to execute the pSUFER protocol on a single protein nanoparticle and analyze the resulting data can be downloaded at the following link: https://files.ipd.uw.edu/pub/JUPITER/JUPITER.tar.gz.

**Figure 2.**
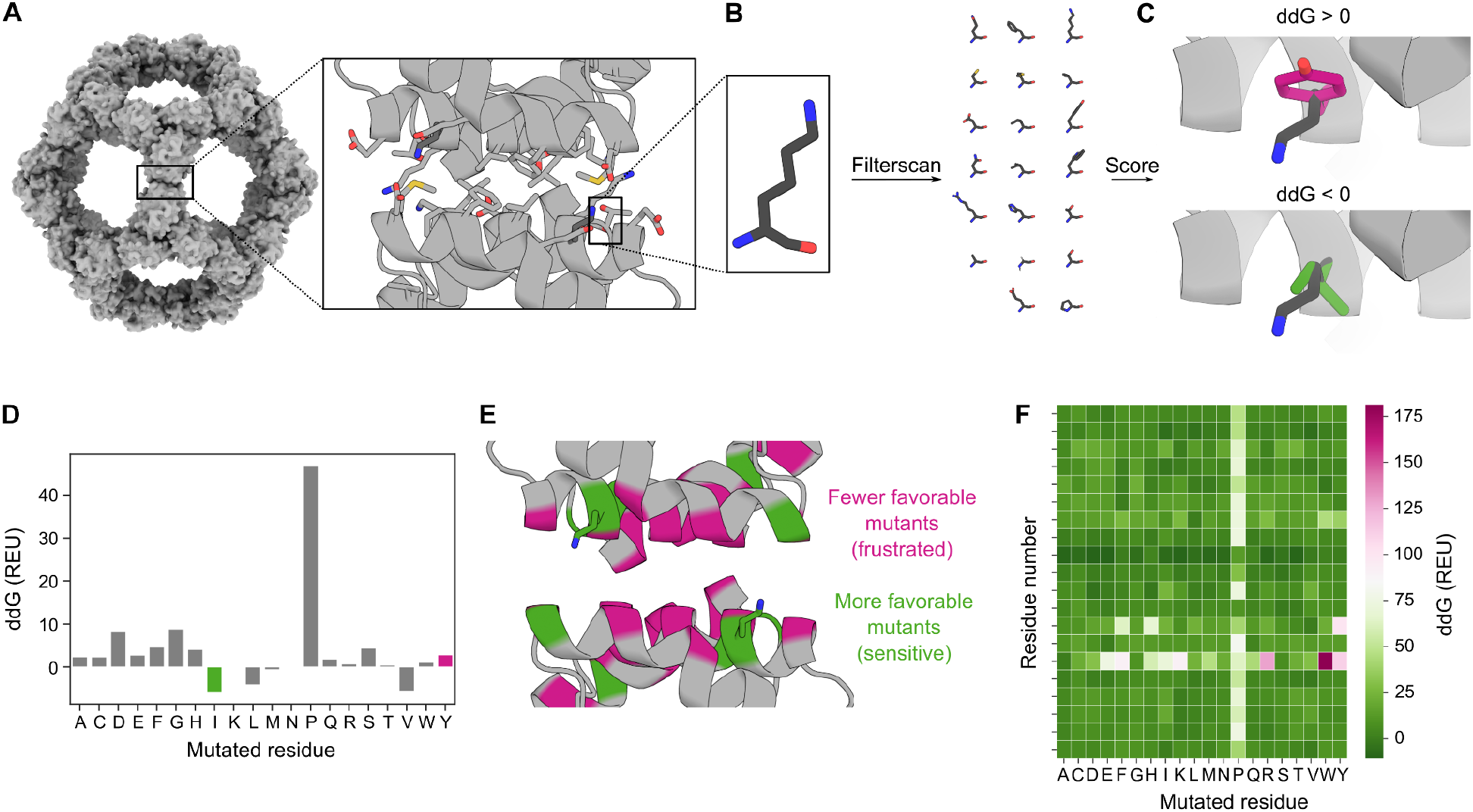
JUPITER’s first phase of research focused on studying individual previously designed successful nanoparticles. Analysis of I3-01^31^ through the pSUFER protocol. **A-C,** pSUFER overview: **A,** The interface between two nanoparticle subunits of a relaxed design model, here with side chains highlighted in stick representation, is selected using ResidueSelectors; **B,** each interface residue (here, K23) is computationally mutated to all possible amino acids; and **C,** the free energy difference compared to the wild-type (ddG) is calculated. **D-F,** Data visualization by the students. **D,** A representative barplot showing the energy difference for each possible mutation at a given sequence position (K32), highlighting a favorable mutation (K23I, green) and an unfavorable mutation (K23Y, pink). **E,** PyMOL visualization of pSUFER scores for the whole interface by differential coloring: *frustrated* residue positions (more than two favorable mutations) are in green while *sensitive* positions (up to two favorable mutations) are in pink. **F,** A heatmap compiling the results: columns represent each possible mutation, and rows represent each interface residue position.

## Phase 2 - Experimental design and results

In phase 2, the undergraduate cohort began to expand the pSUFER protocol they learned in phase 1 to all the nanoparticle designs ever experimentally tested at the Institute for Protein Design between 2012 and July 2020, totalling 809 nanoparticles across 11 icosahedral, octahedral, or tetrahedral architectures. Of these 809 designs, 64 have been experimentally confirmed to adopt the target architecture (“working” designs), while the other 745 apparently failed to assemble (“non-working” designs). Using the pSUFER protocol, the undergraduates generated a single dataset for further analysis, aiming (1) to identify trends that could provide insight into why certain designs may or may not assemble as designed and (2) to predict residue positions where mutations could be introduced to substantially weaken or strengthen protein-protein interfaces across all nanoparticles.

In groups, the students developed computational experiments to probe three hypotheses from a collectively generated pSUFER dataset. One team hypothesized that working nanoparticles would have larger losses in predicted binding energy upon mutation of interface residues compared to non-working nanoparticles (**Figure 3A, left**). This group determined the number of positions at each position of every nanoparticle interface with favorable ddG scores (ddG < 0). Next, the undergraduates compared the fraction of interface positions per nanoparticle that were sensitive or frustrated between working and non-working nanoparticles. They observed no significant difference in the fraction of sensitive and frustrated interface positions between working and non-working nanoparticles (**Figure 3A, right**). Similar analyses were performed on each individual symmetric architecture with a similar outcome.

**Figure 3.**
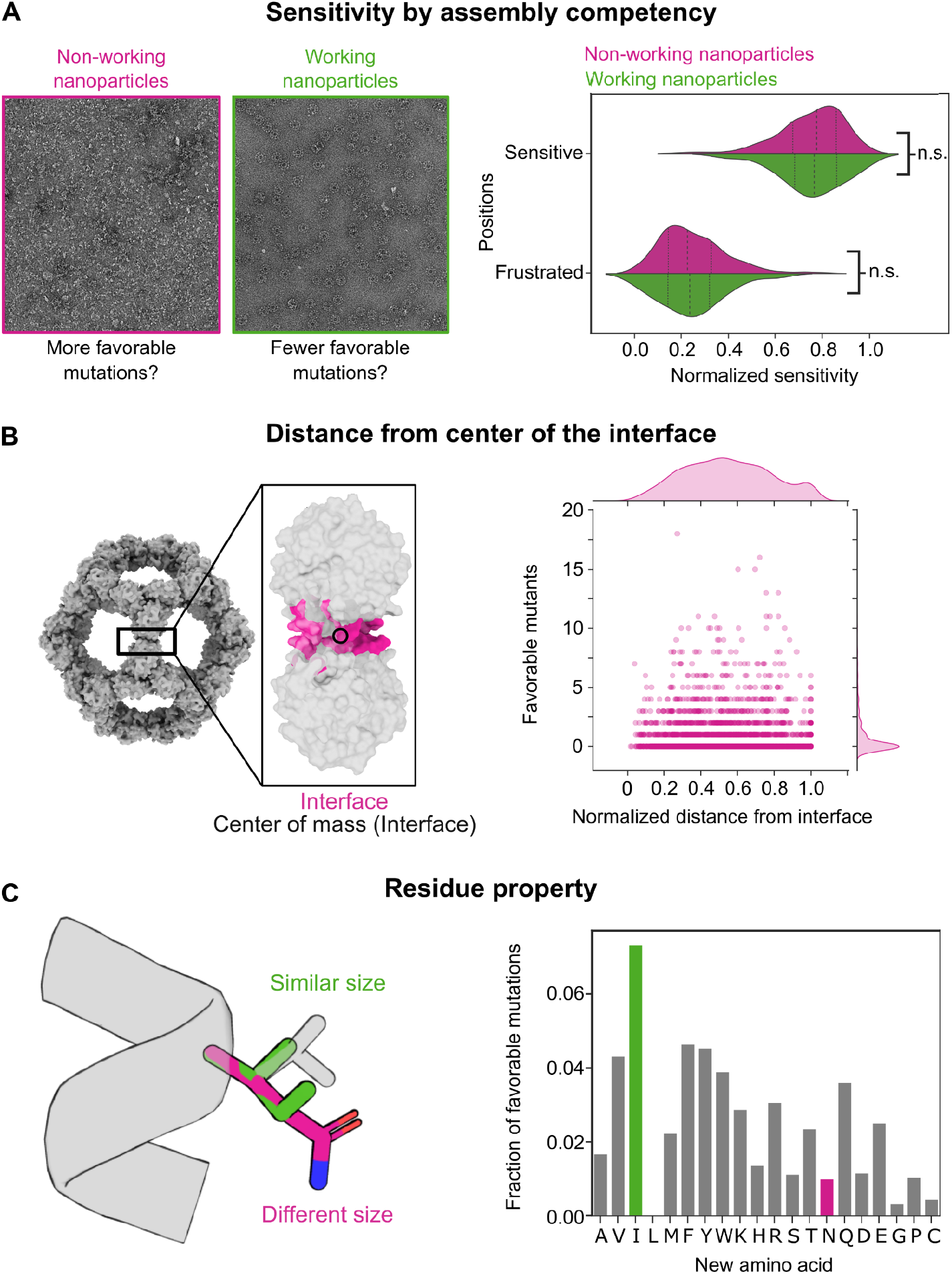
JUPITER’s second phase had students create and test their own hypotheses. **A,** The first group hypothesized that working nanoparticles would be more sensitive (larger predicted losses in Gibbs free energy) to mutations than non-working nanoparticles. However, they observed no significant difference between the mutability ranges of all working vs. all non-working nanoparticles across sensitive, moderate, and frustrated positions (**Supplementary Table 1**). **B,** The second group hypothesized that residues near the center of the interface would be less mutable than those further away; for I3 nanoparticles, they found that positions approximately halfway between the center of mass of the interface and the furthest residue from the interface to be the most mutable. Each circle on the graph represents a single position of a single nanoparticle. **C,** The third group hypothesized that mutations to residues with similar properties (e.g., size, polarity, or charge) would more often be favorable than mutations to residues of differing properties; e.g., for leucines (L) on all nanoparticles, they found that mutations to hydrophobic residues of similar size and characteristics, such as isoleucine (I), were more favorable than to residues introducing larger changes, such as asparagine (N).

The second group hypothesized that positions closer to the center of mass of a nanoparticle interface, by being buried in the designed hydrophobic core, contribute most directly to the protein-protein interface and therefore would be more sensitive (have fewer favorable mutations) compared to positions further from the interface center of mass (**Figure 3B, left**). This group used the pSUFER dataset and the atomic coordinates of each interface position in a nanoparticle to estimate the center of mass of each nanoparticle interface and calculated the number of favorable mutations for each position relative to the position’s Cβ distance from the center of mass of the interface. The group corrected for different interface sizes by normalizing all absolute distances to the largest distance within each nanoparticle interface. Following normalization, the students observed that positions approximately halfway between the center of the interface and the farthest residue from the interface tended to be more frustrated, i.e. to have the most number of favorable mutations per nanoparticle (**Figure 3B, right**).

Finally, the third team hypothesized that amino acid mutations would score more favorably if they were mutated to amino acids with properties—such as size or charge—similar to those of the wild-type residue, as opposed to amino acids with different properties (**Figure 3C, left**). Consistent with this hypothesis, the number of leucine positions with favorable ddG scores upon mutation to isoleucine were higher than the number of leucine positions with favorable ddG scores upon mutation to asparagine (**Figure 3C, right**). Even though leucine, isoleucine, and asparagine are similar in molecular weight, leucine and isoleucine are similar in property as they are both nonpolar residues while asparagine is a polar residue.

**Supplementary Table 1.** Statistical information for comparing position sensitivity by assembly competency. Two-tailed unpaired t-tests were used to compare means (**Figure 3A, right**) with *α* = 0.05 in Graphpad Prism Software.

| Experiment | Condition | n | Mean | Summary | Adjusted p value | t | df |
| --- | --- | --- | --- | --- | --- | --- | --- |
| Sensitive positions | Non-working nanoparticles | 745 | 0.7517 | n.s. | 0.8444 | 0.1963 | 807 |
|  | Working nanoparticles | 64 | 0.7553 | n.s. |  |  |  |
| Frustrated positions | Non-working nanoparticles | 745 | 0.2483 | n.s. | 0.8444 | 0.1963 | 807 |
|  | Working nanoparticles | 64 | 0.2447 | n.s. |  |  |  |

Following our teaching model, the undergraduates could further easily generate three additional datasets to evaluate their hypotheses using simple tweaks to the pSUFER protocol. First, the undergraduates expanded the pSUFER protocol to residues neighboring but not directly participating in the previously analyzed nanoparticle interface (**Figure 4A**). While the number of neighboring residues was lower than that of interface residues, residues in both groups were generally most tolerant of mutations further away from the interface, without many differences observed between groups.

**Figure 4.**
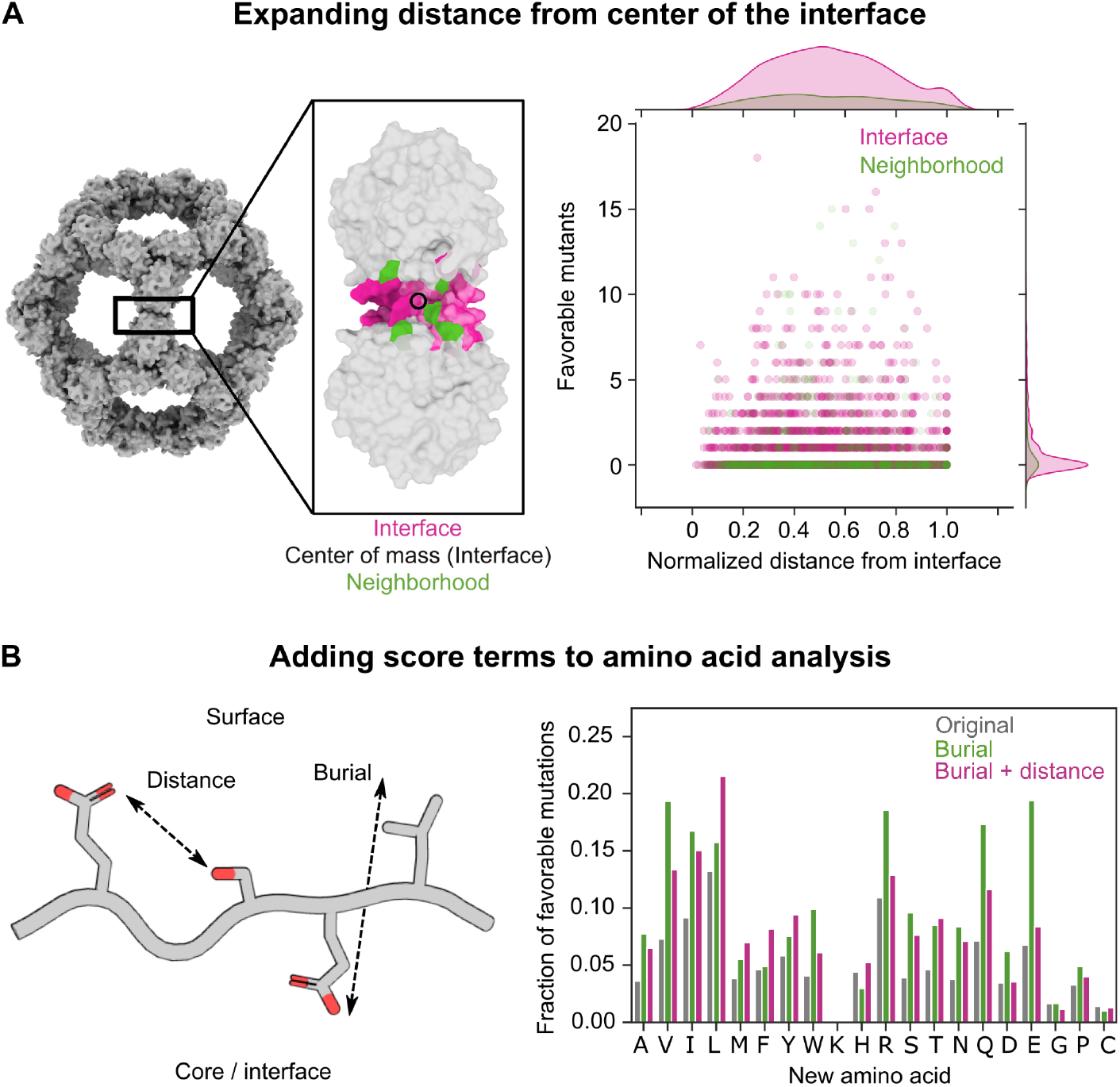
JUPITER students further refined their hypotheses and developed new tests. **A,** Students updated their hypotheses to analyze residues neighboring but not directly participating in the previously analyzed interfaces. The students found that residues neighboring the interface had the most favorable mutations halfway between the center of the mass of the interface and the furthest residue from the interface. Each circle on the graph represents a single position of a single nanoparticle. **B,** The Rosetta score function can be refined by adding distance- and burial-dependent score terms to reward favorable electrostatic interactions (distance) or penalize buried polar residues (burial) based on their depth from the protein surface. By applying two different variants of the score function, the students observed that the addition of burial- and distance-dependent score terms shifted the degree to which different residues lead to favorable mutations across all nanoparticles.

The undergraduates also introduced simple modifications to how ddG scores were evaluated. The score function initially used was an all-atom Rosetta energy score function, dominated by pairwise atomic interactions between protein backbones and side chains, including packing interactions, electrostatic interactions, and implicit solvation^53^. The undergraduates additionally tested pilot burial- and distance-based score terms added to the original score function (*manuscript in preparation*). The burial score term penalized unfavorable polar interactions based on how deeply atoms were buried in the core or interface of the protein, while the distance-based score term rewarded more favorable electrostatic interactions within an optimal distance (**Figure 4B**). The undergraduates added just the burial-dependent score term to generate one dataset and both the burial- and distance-dependent score term to generate the second dataset. By comparing these new datasets to the original one, the undergraduates observed significant differences in the number of favorable mutations across amino acids, especially among polar amino acids.

## Discussion

The JUPITER program successfully re-employed undergraduates who lost research positions due to COVID-19, following a model that could be extended to any research group looking to mentor multiple students at once. The program enabled two to three core advisors to take on a cohort of undergraduate students and invite guests to host one-time lectures and office hours. In this cohort-based learning model, undergraduate researchers can take ownership of one part of a project while working with others to weave their subprojects together into a full story. Our computational study in particular fits well with the remote learning environment imposed by the COVID-19 pandemic. Students successfully learned how to design computational experiments, maintain and analyze large amounts of data in collaboration, and develop scientific hypotheses. Data collected by students is being used to inform future design protocols and improve models that assess protein nanoparticle assembly competency^33^. As hybrid research environments become more popular, the undergraduates from the JUPITER program are returning to research labs, citing the JUPITER program as a major reason for their interest in continuing research.

The student presentations in both phase 1 and 2 provided opportunities to practice scientific communication through both oral presentations and literature review, enabling students to explore the background behind protein nanoparticle design and to develop an understanding of the energetics and kinematics underlying a protein-protein interface. The lecturers observed that the students’ ability to interpret the effects of point mutations at a nanoparticle interface was strengthened as the students developed their own hypotheses and followed up on experiments in phase 2. After the second phase, students reported in a debriefing meeting that their experiences in coding and data interpretation were crucial elements in their desire to further pursue computational research.

The undergraduate researchers provided feedback after both phases of JUPITER through surveys, in which they expressed overwhelming satisfaction. Specifically, out of the 8 responses received on our student surveys, at least 6 students enjoyed each of the weekly lectures in understanding the research topic in phase 1, and 7 students agreed that the JUPITER program gave them a better sense of whether they would like to pursue a career in research. Students expressed that the introductory lectures during phase 1 of the program prepared them well for the large-scale project they carried out in phase 2 and expressed interest in learning more about research topics in protein design. At the end of phase 2, most of the undergraduates expressed appreciation for learning protein design concepts and were excited to learn more about other research projects at our Institute. However, they experienced difficulty studying conceptual topics about protein structure when they were more focused on learning to code and analyze their data, so a future update to the program should provide a more thorough Python or Bash course prior to the start of the research project. The surveys further highlighted the need for good communication and better data management, as well as a desire for undergraduate participants to interact more with researchers outside of the direct JUPITER program. This feedback could inform future cohort-based research programs at our Institute and elsewhere.

During this pilot program, we demonstrated that rigorous research can be achieved in a virtual environment. The undergraduate students learned the necessary coding skills while generating novel scientific datasets that each undergraduate subgroup took into different analysis directions. Such a course format can be applied to any protein system—not only nanoparticles—and nearly any cohort size, and we continue to expand cohort-based mentorship programs in our labs using other protein design protocols. We hope that our model for cohort-based undergraduate research can inspire new research programs across any scientific field, in order to increase the number of highly beneficial research opportunities for aspiring scientists.

## Acknowledgments

We thank Luki Goldschmidt for his instrumental help in getting the JUPITER program undergraduates set up on our remote computing cluster and for maintaining computational resources. We are grateful to Ashley Vater and Justin Siegel for inspiration and advice throughout JUPITER. We thank the Institute for Protein Design and University of Washington Department of Biochemistry staff members Zulfiya Lafi and Lance Stewart for their administrative support. We also thank Dina Listov and Sarel Fleishman for invaluable discussions about the pSUFER protocol. Funding for this work was provided by the Audacious Project at the Institute for Protein Design (R.D., C.S.K., S.C, E.A., Z.S., P.T., H.M., A.F., T.M., J.H., I.O., B.K., C.A., A.K., D.B. and N.P.K.), The Open Philanthropy Project Improving Protein Design Fund (T.F., K.V.W., D.B., N.P.K.), The Nordstrom Barrier Institute for Protein Design Directors Fund (J.O., I.C.H.) NSF DGE-1762114 (E.C.Y.), HDTRA1-19-1-0003 (C.S.K.), the Washington Research Foundation (H.H., K.L.H.), NSF grant CHE-1629214 (N.P.K. and D.B.), and the Howard Hughes Medical Institute (D.B.).

